# Structural basis for loop-dependent inhibition by AcrIIC1 anti-CRISPR proteins

**DOI:** 10.64898/2026.06.11.731568

**Authors:** Shen Wang, Zhiyu Zhang, Chenyu Luan, Cheng Zhu, Chang Liu, Wenling Li, Qianfan Wang, Xiaoqian Liu, Haitao Yang, Yunjie Xiao, Zefang Wang

## Abstract

Most known anti-CRISPR proteins exert inhibition via solitary functional pathways, and few natural Acr homologs have been reported to possess coupled inhibitory mechanisms. Here, we demonstrate that AcrIIC1_Boe_ represses Nme1Cas9 activity through loop-coupled inhibitory effects. Beyond canonical occlusion of the HNH catalytic site, the unique extended loop of AcrIIC1_Boe_ further perturbs sgRNA binding and restricts R-loop maturation to reinforce inhibition. Strikingly, loop deletion completely abrogated this layered inhibitory capacity, confirming that the loop is indispensable for full anti-CRISPR function. Consistently, transplantation of this functional loop into the weakly active AcrIIC1_Nme1_ markedly upgraded its inhibitory performance. Our findings reveal an unprecedented loop-governed coupled inhibition mode, clarify the structural basis for functional divergence within the AcrIIC1 family, and provide a modular engineering strategy for the rational design and optimization of high-potency anti-CRISPR proteins.

## Introduction

Clustered regularly interspaced short palindromic repeats (CRISPR) and CRISPR-associated (Cas) protein systems are adaptive immune systems in bacteria and archaea that defend against reinvasion by mobile genetic elements (MGEs) such as phages and plasmids^1^. These systems confer specific immunity by integrating short segments of foreign DNA into CRISPR loci, which are subsequently transcribed into small guide RNAs that direct Cas effector nucleases to complementary nucleic acid targets for cleavage. Among class II CRISPR systems, Cas9 is the most abundant and diverse effector protein, existing in three subtypes-IIA, IIC, and the relatively rare IIB-of which types IIA and IIC are the most prevalent^2^. The programmable nature of Cas9 has made it a powerful tool for gene editing and genomic manipulation across a wide range of organisms, yet its native biological function remains subject to intense selective pressure from phage-encoded inhibitors^3^.

In response, phages and other MGEs have evolved Anti-CRISPR proteins (Acrs) that inhibit CRISPR-Cas function through a remarkably diverse array of strategies^4^. Acrs are typically small proteins that share limited sequence or structural homology with known protein families, reflecting their rapid evolution under strong selective pressure. The inhibitory mechanisms characterized to date span nearly every step of the CRISPR-Cas immune response^5^. Anti-CRISPR proteins deploy highly diverse molecular toolkits to paralyze host CRISPR-Cas machineries at multiple regulatory inflection points. Ribonucleoprotein (RNP) complex assembly can be intercepted at the onset by factors like AcrIIA16-AcrIIA19, AcrIIC2, and AcrVA4^6–8^, which bind Cas9 and sterically hinder single guide RNA (sgRNA) engagement, a mechanism mirrored by the Cas13a-inhibitor AcrVIA1 during substrate RNA selection^9^. Alternatively, mature complexes can be prevented from executing nucleic acid hybridization. This is achieved by subunit interference with crRNA-DNA pairing (as seen with AcrIF1, AcrIF8, AcrIF9, and AcrIF14)^10–13^, or structural deception using charge-surface DNA mimics to mask PAM recognition sites (including AcrIIA2, AcrIIA4, AcrIIA15, AcrIIC5, and AcrVA1)^14–17^. Beyond direct site occlusion, long-range allosteric modulation represents an elegant pathway, wherein regulators like AcrIIA6, AcrII25.1, and AcrIIA32 dynamically lower Cas9’s target affinity by destabilizing its structural elasticit^18,19^. These classical paradigms are further supplemented by unconventional stratagems, such as regulatory post-translational modifications or the chemical interception of chemical second messengers by Type III Acr variants. More recently, AcrIIA27 was shown to prevent R-loop formation^20^.

Among the characterized Acrs are those targeting type II CRISPR-Cas9 systems, several of which inhibit Cas9 by binding directly to the conserved HNH nuclease domain and occluding its catalytic site. Within this group, the AcrIIC1 family has emerged as a conserved lineage of broad-spectrum inhibitors targeting type II-C Cas9 orthologs. The founding member, AcrIIC1_Nme1_, is an 85-amino-acid protein that binds the HNH domain of Nme1Cas9 and traps it in a catalytically inactive state by forming hydrogen bonds with the catalytic residues H588 and D587, effectively converting Cas9 into a dCas9^21–23^. This HNH-occlusion mechanism has served as a paradigm for understanding AcrIIC1 function, and structural studies have revealed that the AcrIIC1-HNH binding interface centers on a conserved set of residues that directly engage the catalytic center.

Despite the mechanistic diversity of Acr proteins, most known Acr proteins exert their inhibitory effects through solitary pathways, and coupled inhibitory mechanisms in natural Acr homologs have rarely been reported. This knowledge gap limits our understanding of the functional diversity and regulatory complexity of Acr proteins. Here, we show that AcrIIC1_Boe_ employs an extended loop region that critically contributes to Nme1Cas9 inhibition. While AcrIIC1_Boe_ retains the conserved HNH catalytic-site interaction network characteristic of the AcrIIC1 family, its loop region additionally approaches the sgRNA spacer-binding region adjacent to the HNH domain and influences Nme1Cas9 interactions with guide RNA and target DNA. Through structural, biochemical, and biophysical analyses, we demonstrate that the loop region is broadly required for productive AcrIIC1-HNH engagement and that transplantation of the AcrIIC1_Boe_ loop into AcrIIC1_Nme1_ enhances inhibitory activity. Collectively, our findings uncover a loop-governed coupled inhibition mode, elucidate the functional differences within the AcrIIC1 family, and provide valuable insights for the rational modular optimization of potent Anti-CRISPR proteins.

## Results

### AcrIIC1_Boe_ exhibits enhanced inhibitory activity despite conserved HNH engagement

Firstly, we compared the inhibitory effects of AcrIIC1_Boe_ and AcrIIC1_Nme1_ on Nme1Cas9-mediated target DNA cleavage in vitro at varying concentrations (**Fig. S1**). As illustrated in **Fig. 1A**, AcrIIC1_Boe_ fully suppressed Nme1Cas9 activity at a molar ratio of 2:1 relative to Nme1Cas9. In contrast, AcrIIC1_Nme1_ exerted complete inhibition only at 4-fold molar excess. This discrepancy demonstrates that AcrIIC1_Boe_ possesses superior inhibitory potency.

**Figure 1.**
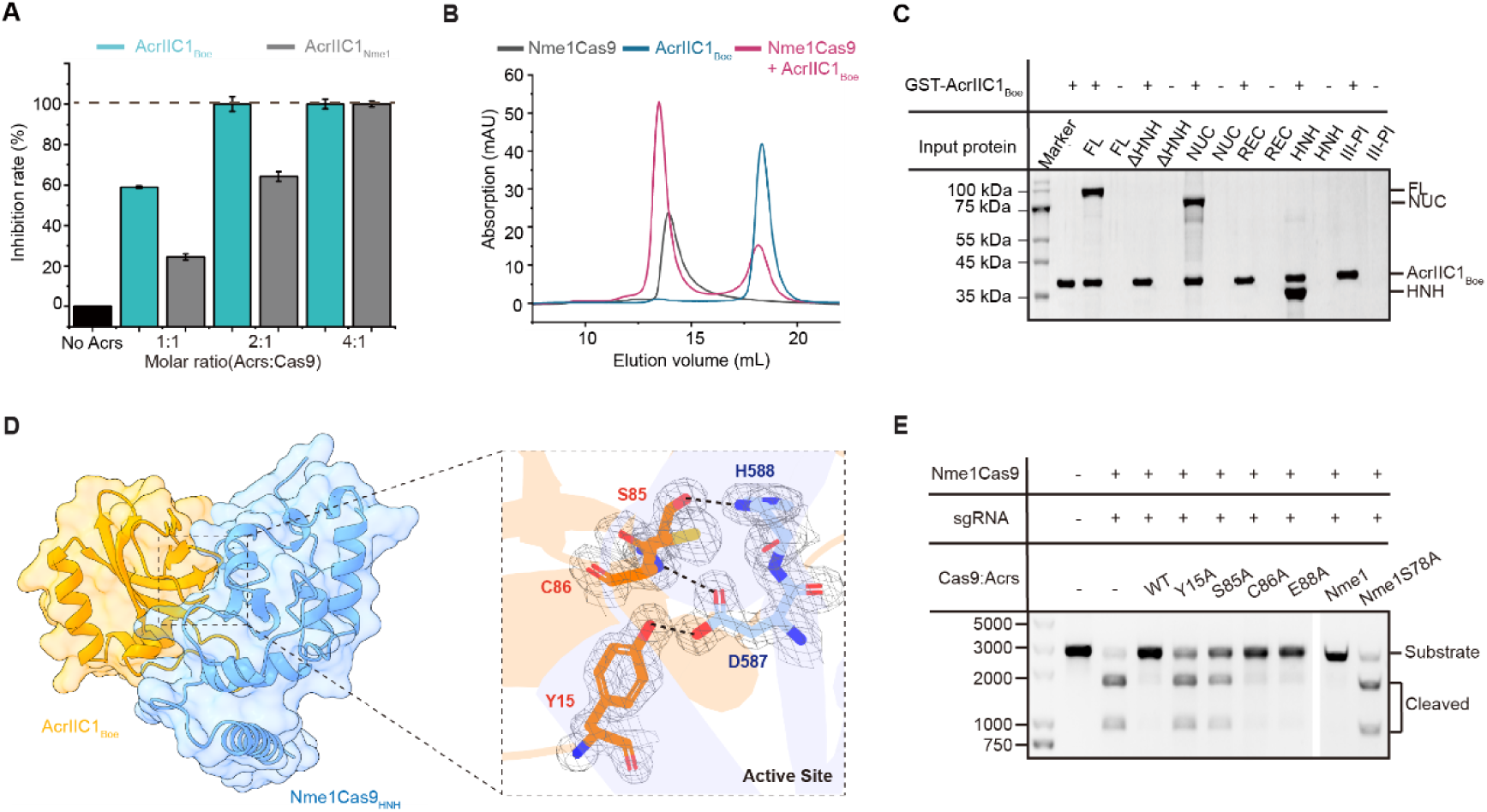
AcrIIC1_Boe_ inhibits Nme1Cas9 more potently than AcrIIC1_Nme1_ despite forming an identical HNH binding interface. **A,** DNA cleavage assays performed with Nme1Cas9 in the presence of AcrIIC1_Boe_ and AcrIIC1_Nme1_ at various ratios. **B,** Nme1Cas9, AcrIIC1_Boe_, and the Nme1Cas9-AcrIIC1_Boe_ complex were eluted via size-exclusion chromatography using Superdex 200 increase. **C,** Pull-down results of AcrIIC1_Boe_ with Nme1Cas9 and truncated domains of Nme1Cas9. **D,** Occlusion of HNH active site residues (blue) through hydrogen bonding with AcrIIC1_Boe_ (orange). HNH catalytic residues H588 and D587 form hydrogen bonds (black dotted line) with Y15, S85 and C86 of AcrIIC1_Boe_, respectively. 2mFo-DFc electron density map is shown for interacting residues and contoured at 1.6 σ. **E,** DNA cleavage assays conducted by Nme1Cas9 in the presence of WT AcrIIC1 and the indicated mutants. Each experiment was performed in triplicate (n=3). Statistical analysis was performed using a one-way ANOVA, followed by post-hoc Tukey’s test for multiple comparisons. Data are presented as mean ± SEM.

Previous studies revealed that AcrIIC1_Nme1_ inhibits Nme1Cas9 by associating with its HNH domain and masking key catalytic residues D587 and H587 responsible for DNA cleavage^21^. To explore whether the divergent inhibitory potency stems from distinct binding modes toward Nme1Cas9, we first verified the interaction between AcrIIC1_Boe_ and Nme1Cas9 via size-exclusion chromatography (**Fig. 1B**). The elution peak of mixed protein samples emerged earlier than that of isolated Nme1Cas9, confirming stable complex formation.

To pinpoint the exact binding region, we constructed a series of Nme1Cas9 domain-truncated mutants (**Fig. S2**)^24^. Pull-down assays showed that AcrIIC1_Boe_ interacted with full-length Nme1Cas9, NUC variant and standalone HNH domain, yet failed to bind ΔHNH constructs. These findings demonstrate both AcrIIC1_Boe_ and AcrIIC1_Nme1_ target the HNH domain of Nme1Cas (**Fig. 1C**)^25,26^.

To further characterize the binding mode, we solved the crystal structure of AcrIIC1_Boe_-HNH complex at 1.8 Å resolution. Structural observation revealed tight binding of AcrIIC1_Boe_ at the catalytic pocket of HNH domain (**Fig. 1D, left**). Structural alignment with the published AcrIIC1_Nme1_-HNH complex (PDB: 5VGB) yielded an RMSD of 0.688 Å across 1154 aligned atoms (**Fig. S3**). The low RMSD value reflects high structural similarity between two complexes, implying that binding mode difference is not the cause of their potency discrepancy.

At the interaction interface, catalytic residues D587 and H588 of Nme1Cas9 form hydrogen bonds with Y15, S85 and C86 of AcrIIC1_Boe_ (**Fig. 1D, right**), which is highly consistent with the binding pattern of AcrIIC1_Nme1_. Extra hydrogen bonds including E88-N616, E88-S593 and S85-F592 further stabilize the intermolecular interface (**Fig. S4**).

Alanine scanning mutagenesis was performed to evaluate the functional significance of interfacial residues. The Y15A mutation diminished inhibitory activity by nearly 90%, whereas C86A and E88A mutants retained wild-type potency, consistent with previous AcrIIC1_Nme1_ research. Notably, S85A mutant maintained half of the original inhibitory capacity, while the equivalent S78A substitution in AcrIIC1_Nme1_ almost abolished its suppression effect^21^.

Collectively, AcrIIC1_Boe_ and AcrIIC1_Nme1_ share highly conserved binding modes when engaging the HNH domain. Such conserved interaction cannot rationalize the stronger inhibitory capacity of AcrIIC1_Boe_, indicating the existence of an extra regulatory mechanism underlying its enhanced suppression effect.

### AcrIIC1_Boe_ loop affects sgRNA spacer positioning through HNH-associated interactions

To further investigate the molecular mechanism underlying the stronger inhibitory activity of AcrIIC1_Boe_, we conducted 100 ns molecular dynamics (MD) simulations of the Nme1Cas9-sgRNA complex under different Acr protein conditions (**Fig. 2A and S5**)^27^. Trajectory analysis revealed that the loop region of AcrIIC1_Boe_ inserts into the concave groove of the Nme1Cas9 HNH domain, a structural pocket that accommodates the 5′ segment of the sgRNA spacer (**Fig. 2B and S6**)^27^. In contrast, the truncated and shorter loop region of AcrIIC1_Nme1_ cannot reach this functional groove and thus produces no steric hindrance (**Fig. S7**). These observations indicate that the AcrIIC1_Boe_ loop may perturb the stable docking of the sgRNA spacer on the HNH domain. Consistent with this potential structural interference, root mean square fluctuation (RMSF) analysis of the first six nucleotides of the sgRNA spacer throughout the 100 ns simulations showed markedly enhanced conformational flexibility in the AcrIIC1_Boe_-bound system compared with the Acr-free control group (**Fig. 2C**).

**Figure 2.**
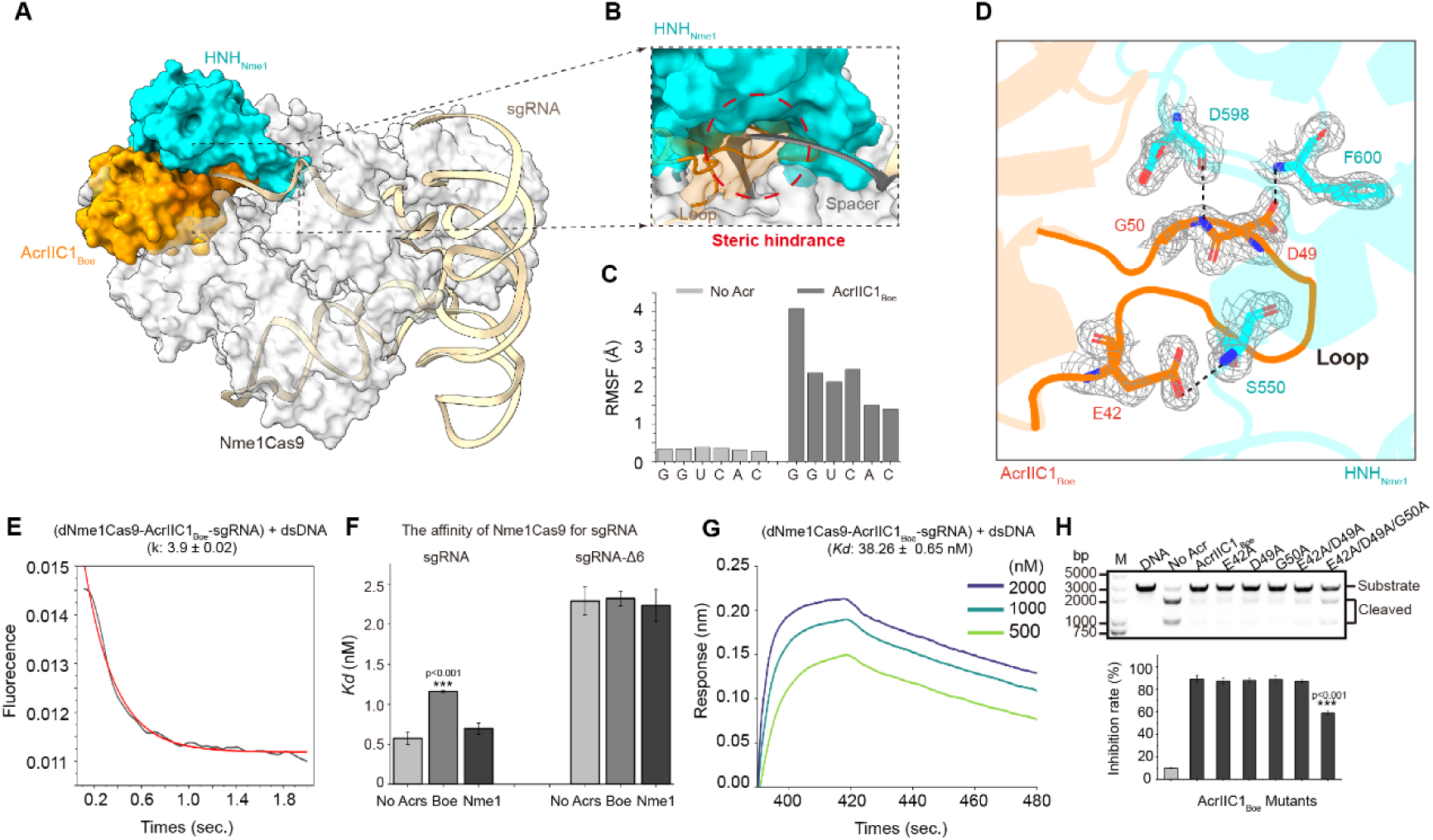
The AcrIIC1_Boe_ loop interferes with sgRNA spacer positioning and affects R-loop formation. **A,** Molecular dynamics simulation of the Nme1Cas9-sgRNA complex in the presence of AcrIIC1_Boe_. AcrIIC1_Boe_ is shown in orange, the HNH domain in cyan, and sgRNA in wheat. **B,** Enlarged view showing overlap between the AcrIIC1_Boe_ loop and the 5′ end of the sgRNA spacer region. **C,** Root-mean-square fluctuation (RMSF) analysis of the first six nucleotides of the sgRNA spacer during 100-ns molecular dynamics simulations in the absence or presence of AcrIIC1_Boe_. **D,** Hydrogen-bond interactions between the AcrIIC1_Boe_ loop and the HNH domain observed in the AcrIIC1_Boe_-HNH crystal structure. Hydrogen bonds are indicated by dashed lines. **E,** R-loop formation kinetics measured by stopped-flow FRET in the presence of AcrIIC1_Boe_. **F,** Binding affinity measurements between Nme1Cas9 and full-length sgRNA or Δ6-sgRNA in the absence or presence of Acr proteins. **G,** Binding affinity measurements between dNme1Cas9-sgRNA and dsDNA in the presence of AcrIIC1_Boe_. **H,** In vitro DNA cleavage assays of AcrIIC1_Boe_ loop mutants. Quantification of cleavage inhibition is shown below. Data are presented as mean ± SEM from three independent experiments.

To clarify why the AcrIIC1_Boe_ loop can maintain such a specific binding posture, we analyzed the hydrogen bond network between the loop and the HNH domain and discovered multiple interactions which have not been documented in previous related study^21^. As shown in **Fig. 2D**, three residues in the AcrIIC1_Boe_ protein (E42, D49, G50) formed hydrogen bonds with the HNH domain (S550, F600, and D598). These structural interactions may help maintain the orientation of the loop observed near the sgRNA Spacer, thereby exerting spatial steric hindrance on the 5’ end of the sgRNA spacer region. It is noteworthy that the interaction between G50 and D598 in the loop region led to a conformational change in D598, enabling it to form a hydrogen bond with the catalytic residue H588 (**Fig. S8**). In contrast, AcrIIC1_Nme1_ only binds to H588 through S78, which explains why, after the S85 mutation in AcrIIC1_Boe_, the H588 in the HNH domain was still bound by the special conformation of the AcrIIC1_Boe_ loop region, thus retaining a certain inhibitory activity (**Fig. 1E**).

Given the structural evidence that the anchored loop disturbs the spatial arrangement of sgRNA spacer, we next test whether the AcrIIC1_Boe_ loop affects the process of Nme1Cas9 R-loop formation by using the stopped-flow FRET method to measure the kinetics of R-loop formation. Without Acr, the observed rate constant for R-loop formation (k_obs_) was 8.4 ± 0.06 s⁻¹, while for AcrIIC1_Nme1_ it was 8.0 ± 0.19 s⁻¹. In contrast, AcrIIC1_Boe_ reduced the rate constant to 3.9 ± 0.02 s⁻¹ (**Fig. 2E and S9**). These results indicate that AcrIIC1_Boe_ slows down R-loop formation rather than completely abolishing this process^28^.

Next, we investigated how this loop structure affects the interaction between Nme1Cas9-sgRNA. The bioluminescence imaging (BLI) measurements showed that the dissociation constant of the binding of Nme1Cas9 to the full-length sgRNA was 0.58 ± 0.07 nM (without Acr), 0.70 ± 0.07 nM (AcrIIC1_Nme1_), and 1.16 ± 0.01 nM (AcrIIC1_Boe_) (**Fig. 2F**)^20^. Using the truncated sgRNA (Δ6-sgRNA) that lacks the first six nucleotides of the spacer region, the dissociation constants in all conditions were similar (∼2.2 nM), and AcrIIC1_Boe_ lost its effect on the Nme1Cas9-sgRNA interaction, which confirmed that the AcrIIC1_Boe_ loop is more likely to affect the interaction involving the 5’ end of the spacer region, which is consistent with the results of molecular dynamics (MD) and R-loop analysis^29^. Size exclusion chromatography confirmed that the ternary Nme1Cas9-AcrIIC1_Boe_-sgRNA complex still formed (**Fig. S10**), suggesting the loop does not block sgRNA binding. Limited proteolysis assays showed intensified protein bands of the Cas9-sgRNA complex upon AcrIIC1_Boe_ addition, reflecting elevated conformational flexibility of the complex (**Fig. S11**)^30,31^.

To assess the interference of the loop on the Spacer region and further investigate the downstream effects on the interaction between Nme1Cas9-sgRNA-DNA, we first measured the binding of the dNme1Cas9-sgRNA complex to the target double-stranded DNA (dsDNA). The EMSA results showed that without sgRNA, dNme1Cas9 could not bind to the dsDNA. The ternary dNme1Cas9-sgRNA complex bound to the dsDNA at 160 nM. In the presence of AcrIIC1_Nme1_, binding was also detected at 160 nM, while AcrIIC1_Boe_ increased the concentration required for detectable binding to 220 nM (**Fig. S12**)^32,33^. Meanwhile, consistent BLI measurements gave *Kd* values of 22.77 ± 1.28 nM, 24.09 ± 0.49 nM, and 38.26 ± 0.65 nM for the control, AcrIIC1_Nme1_ and AcrIIC1_Boe_ groups respectively (**Fig. 2F and S13**)^20^. These findings confirm AcrIIC1_Boe_ weakens Cas9-sgRNA-DNA interaction, matching the suppressed R-loop formation kinetics.

Finally, in order to directly test the contribution of the loop-HNH domain interaction to the enhancement of inhibitory effect, we generated a triple mutant of AcrIIC1_Boe_ (E42A/D49A/G50A) which eliminated all three hydrogen bonds. The mutant exhibited approximately 40% reduction in inhibitory activity (**Fig. 2H**). Pull-down assays still detected stable binding between the mutant and HNH domain (**Fig. S14**), indicating disruption of the hydrogen bond network impairs proper loop positioning required for spacer interference, and consequently attenuates suppression efficiency. These results indicate that the AcrIIC1_Boe_ loop region, which is stabilized by specific hydrogen bonds formed with the HNH domain, is located near the sgRNA spacer region. This positional relationship slows down the formation of the R-loop, reduces the affinity of Nme1Cas9 for the sgRNA and target DNA, and provides a cooperative mechanism independent of the HNH active site closure.

### The Loop region is essential for AcrIIC1-Cas9 binding and inhibitory function

Based on the functional role of the AcrIIC1_Boe_ loop region in the interference and enhancement inhibition effects of Spacer, we next investigated whether this loop region is a necessary component of the inhibitory effect mediated by AcrIIC1. To address this question, we constructed loop-deletion mutants of AcrIIC1_Boe_ and AcrIIC1_Nme1_, designated AcrIIC1_Boe_-Δloop and AcrIIC1_Nme1_-Δloop (**Fig. S15**). In vitro DNA cleavage assays were performed to evaluate their inhibitory capability. Surprisingly, as shown in **Fig. 3A**, loop truncation completely abolished the inhibitory activity of both proteins. To uncover the mechanism underlying activity loss, we carried out pull-down assays using the isolated HNH domain. Both loop-deletion mutants exhibited markedly weakened binding affinity toward the HNH domain relative to their wild-type counterparts (**Fig. 3B**), demonstrating that the loop facilitates stable association between AcrIIC1 and the HNH domain.

**Figure 3.**
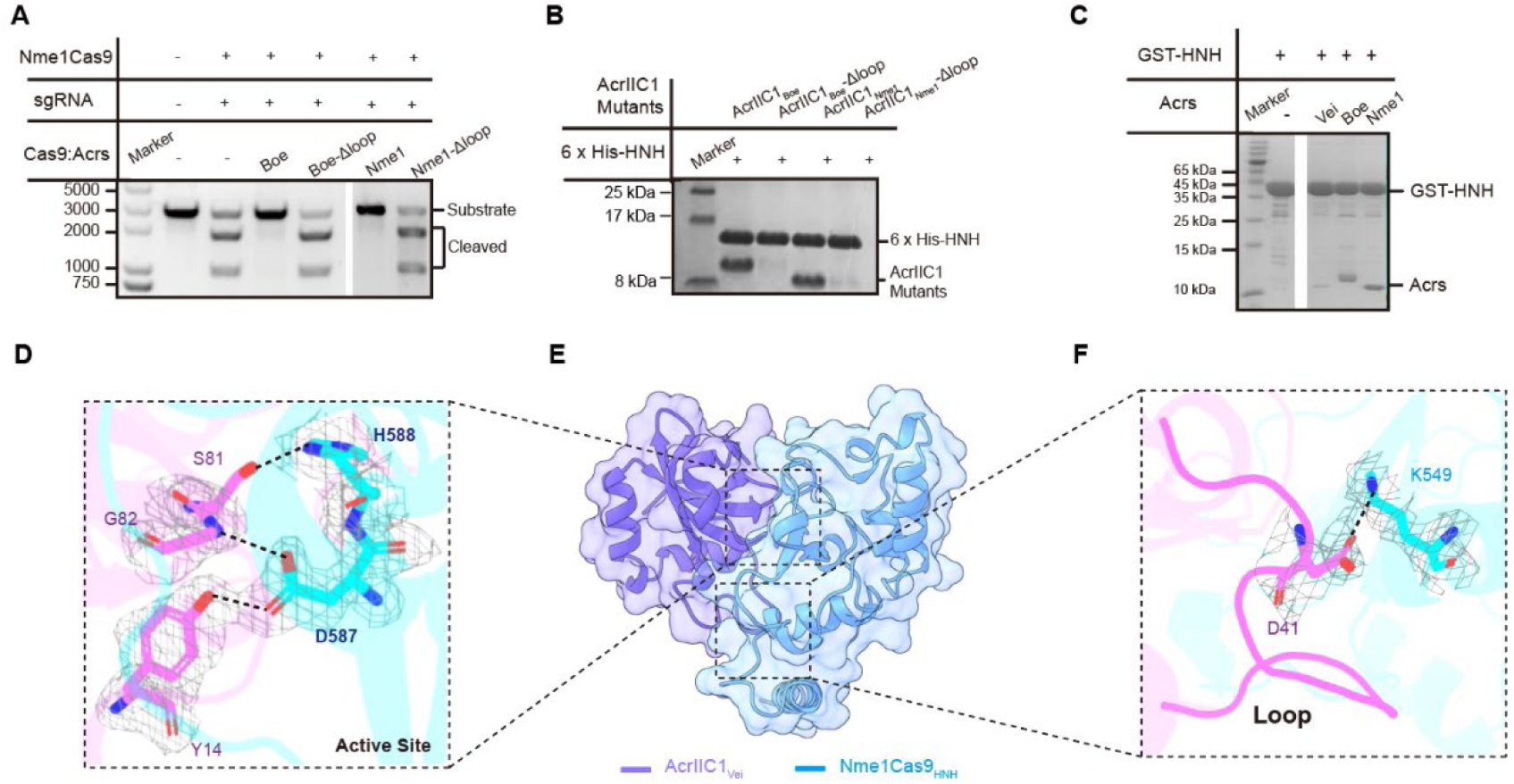
The loop region is required for effective AcrIIC1 inhibition. **A,** In vitro DNA cleavage assays of AcrIIC1_Boe_ and AcrIIC1_Nme1_ loop-deletion mutants. **B,** Pull-down assays of AcrIIC1_Boe_ and AcrIIC1_Nme1_ loop-deletion mutants with the isolated HNH domain. **C,** Pull-down assays comparing interactions of AcrIIC1_Vei_, AcrIIC1_Boe_, and AcrIIC1_Nme1_ with the isolated HNH domain. **D,** Close-up view of the catalytic-site interaction network in the AcrIIC1_Vei_-HNH complex structure. Electron density maps are shown as gray mesh. **E,** Overall crystal structure of the AcrIIC1_Vei_-HNH complex. AcrIIC1_Vei_ is shown in purple and the HNH domain in cyan. **F,** Close-up view of the AcrIIC1_Vei_ loop region interacting with the HNH surface. Electron density maps are shown as gray mesh.

To further elucidate the structural basis underlying the functional divergence of AcrIIC1 family members, we characterized a naturally inactive AcrIIC1_Vei_ homolog newly identified in our study. Consistent with our in vitro cleavage assays, AcrIIC1_Vei_ exhibited no inhibitory activity against Nme1Cas9 (**Fig. S16**). Although structural prediction and sequence alignment revealed conserved core residues (Y14 and S81) and overall structural similarity with active AcrIIC1 homologs, AcrIIC1_Vei_ displayed distinct loop length and positional features (**Fig. S17**). Functionally, pull-down assays confirmed that AcrIIC1_Vei_ possessed extremely weak binding affinity for the HNH domain compared with active AcrIIC1_Boe_ and AcrIIC1_Nme1_ (**Fig. 3C**). To precisely resolve the structural cause of its impaired function, we solved the crystal structure of the AcrIIC1_Vei_-HNH complex (**Fig. 3E**). Structural analysis showed that AcrIIC1_Vei_ retains the ability to occupy the HNH catalytic surface, where its Y14 and S81 form hydrogen bonds with the catalytic residues H588 and D587. However, the divergent loop region of AcrIIC1_Vei_ only mediates a single weak hydrogen bond between D41 and HNH K549, with no additional stable intermolecular interactions observed (**Fig. 3F**). This minimal loop-mediated interaction fails to support stable protein docking.

Overall, these research results indicate that the loop region is responsible for enabling the AcrIIC1 protein to be correctly localized on the surface of HNH during the formation of the complex. The disruption or weakening of these interactions mediated by the loop region will reduce the binding degree of HNH, thereby preventing the AcrIIC1 protein from forming a functional inhibitory conformation and eliminating its inhibitory activity.

### The loop region regulates AcrIIC1 inhibitory activity

Our above results demonstrate that the loop region is a core functional module governing the inhibitory potency of AcrIIC1 proteins, and the unique structural arrangement of the AcrIIC1_Boe_ loop substantially enhances its anti-CRISPR activity. To further verify whether this loop module is transferable and determines functional divergence among AcrIIC1 homologs, we performed loop-swapping experiments between AcrIIC1_Boe_ and AcrIIC1_Nme1_, generating two chimeric mutants, AcrIIC1_BoeST_ and AcrIIC1_Nme1ST_ (**Fig. 4A**). In vitro DNA cleavage assays showed that AcrIIC1_Boe_ completely lost its inhibitory capability toward Nme1Cas9 after grafting the AcrIIC1_Nme1_ loop (**Fig. 4B and 4C**). In contrast, replacing the endogenous loop of AcrIIC1_Nme1_ with the AcrIIC1_Boe_ loop significantly improved its inhibitory efficiency. Wild-type AcrIIC1_Nme1_ required a four-fold molar excess relative to Nme1Cas9 for complete inhibition, whereas AcrIIC1_Nme1ST_ achieved equivalent inhibition at only a two-fold excess, exhibiting an inhibitory level comparable to wild-type AcrIIC1_Boe_.

**Figure 4.**
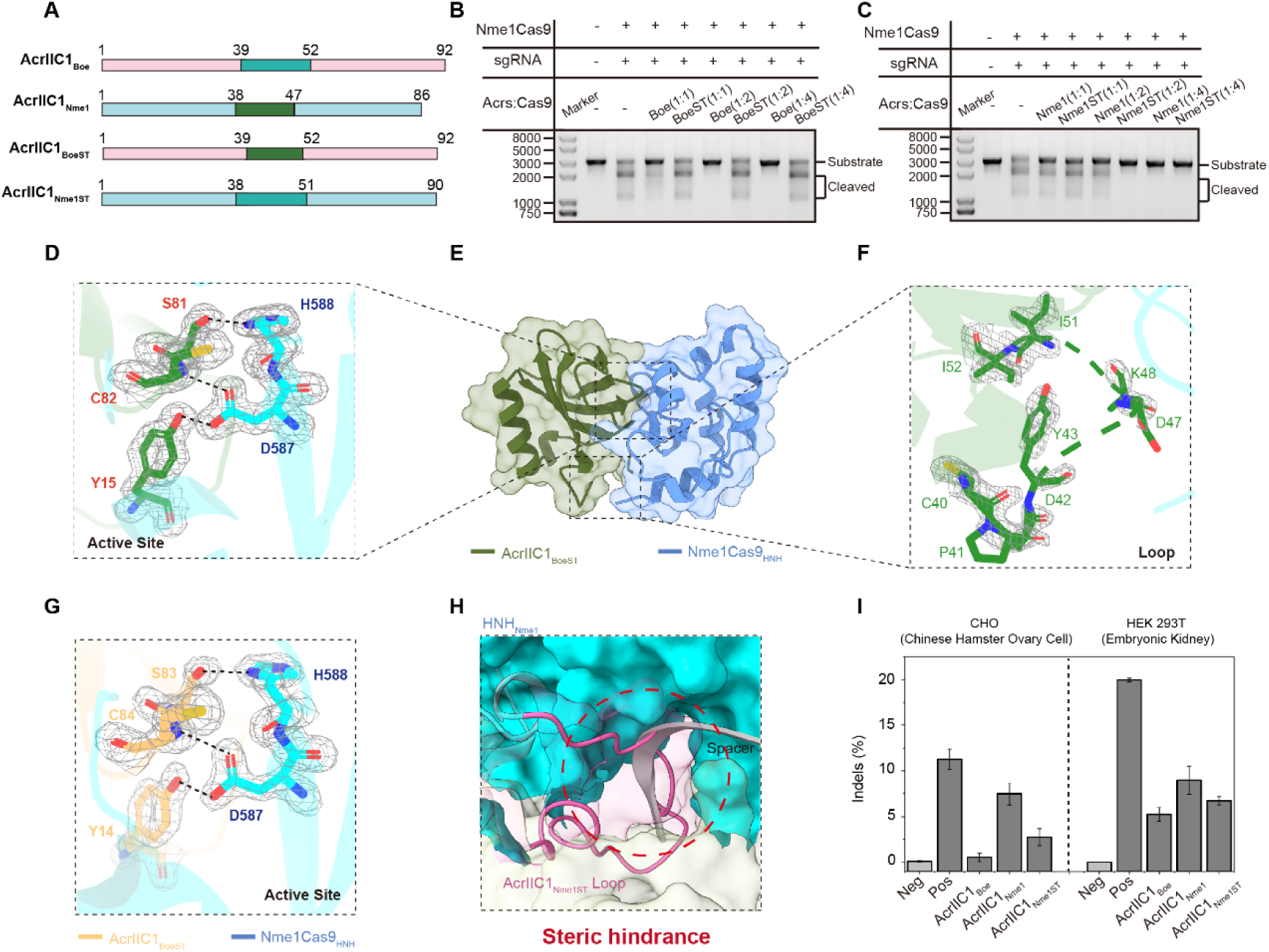
Loop transplantation confers sgRNA interference activity and enhances the inhibitory potency of AcrIIC1_Nme1_. **A,** Schematic diagram of the molecular construction of AcrIIC1_BoeST_ and AcrIIC1_Nme1ST_. **B,** In vitro DNA cleavage assays of AcrIIC1_Boe_ and AcrIIC1_BoeST_ at various ratios. **C,** In vitro DNA cleavage assays of AcrIIC1_Nme1_ and AcrIIC1_Nme1ST_ at various ratios. **D,** Close-up view of the catalytic-site interaction network in the AcrIIC1_BoeST_-HNH complex structure. Electron density maps are shown as gray mesh. **E,** Overall crystal structure of the AcrIIC1_BoeST_-HNH complex. AcrIIC1_BoeST_ is shown in green and the HNH domain in cyan. **F,** Close-up view of the AcrIIC1_BoeST_ loop region interacting with the HNH surface. Electron density maps are shown as gray mesh. **G,** Close-up view of the catalytic-site interaction network in the AcrIIC1_Nme1ST_-HNH complex structure. Electron density maps are shown as gray mesh. **H,** Enlarged view showing overlap between the AcrIIC1_Nme1ST_ loop and the 5′ end of the sgRNA spacer region. **I,** TIDE analysis of the inhibitory efficiency of AcrIIC1_Boe_, AcrIIC1_Nme1_ and AcrIIC1_Nme1ST_ on Nme1Cas9 in CHO cells and HEK293T cells. Each experiment was performed in triplicate (n=3). Statistical analysis was performed using a one-way ANOVA, followed by post-hoc Tukey’s test for multiple comparisons. Data are presented as mean ± SEM.

To uncover the structural basis for the distinct functional outcomes of loop swapping, we first solved the crystal structure of the AcrIIC1_BoeST_-HNH complex (**Fig. 4E**). Structural analysis showed that the interaction network at the catalytic site remained largely stable. Specifically, residues Y15, C82, and S81 in AcrIIC1_BoeST_ still formed hydrogen bonds with the catalytic residues H588 and D587 in the HNH domain (**Fig. 4D**), indicating that occlusion of the active site was not compromised. However, loop substitution markedly increased the conformational flexibility of the AcrIIC1_BoeST_ loop region, resulting in incomplete and discontinuous electron density that precluded full structural resolution (**Fig. 4F**). This observation indicates that heterologous loop replacement disrupts the stable and fixed loop conformation required for efficient inhibition.

We next determined the crystal structure of the AcrIIC1_Nme1ST_-HNH complex to elucidate the structural mechanism underlying its enhanced activity (**Fig. S18**). Structural comparison with wild-type AcrIIC1_Nme1_-HNH revealed a highly conserved overall binding mode and stable catalytic interface. Residues Y14, S83, and C84 of AcrIIC1_Nme1ST_ retained hydrogen-bonding interactions with HNH H588 and D587, maintaining intact active-site shielding (**Fig. 4G**).

To determine whether the enhanced inhibitory activity of AcrIIC1_Nme1ST_ is due to the spacer interference characteristic observed in AcrIIC1_Boe_, we conducted molecular dynamics simulations on the Nme1Cas9-sgRNA complex in the presence of AcrIIC1_Nme1ST_. Similar to AcrIIC1_Boe_, the transplanted loop was close to the sgRNA spacer region and overlapped with the distal spacer nucleotide part of the PAM (**Fig. 4H**), suggesting that the loop region of AcrIIC1_Nme1ST_ also interfered with the spacer.

We further validated these in vitro observations in mammalian cell lines (HEK293T and CHO) via TIDE assays^34^. Consistent with the in vitro biochemical data, AcrIIC1_Boe_ exhibited the strongest inhibitory effect, while the activity of AcrIIC1_Nme1_ was the weakest. In both cell types, the inhibitory efficiency of AcrIIC1_Nme1ST_ was between these two proteins (**Fig. 4I**), confirming that loop transplantation robustly improves AcrIIC1 inhibitory performance in a cellular context.

Subsequently, we further evaluated whether transplanting the AcrIIC1_Boe_ loop into AcrIIC1_Nme1_ would alter the inhibitory profile of AcrIIC1_Nme1_. However, compared to the wild-type AcrIIC1_Nme1_, AcrIIC1_Nme1ST_ did not show enhanced inhibitory activity towards other Cas9 homologs (including CjeCas9, GeoCas9, SauCas9, and SpyCas9) (**Fig. S19**)^21^. These results indicate that the loop-mediated activity enhancement is strictly dependent on the Nme1Cas9 structural microenvironment and does not confer broad-spectrum anti-CRISPR activity.

Finally, in order to further investigate whether the loop exchange technology is compatible with the additional AcrIIC1 engineering strategy, we constructed variants of AcrIIC1_Nme1_ and AcrIIC1_Nme1ST_ that were fused with mCherry and evaluated their inhibitory activities in vitro^35^. As shown in **Fig. S20**, when the molar concentration ratio of Acr to Nme1 Cas9 was the same, the inhibitory activity of AcrIIC1_Nme1_ was the lowest. Both AcrIIC1_Nme1ST_ and AcrIIC1_Nme1_-mCherry improved their inhibitory activity to a certain extent, and when combined, the inhibitory activity reached the highest level. These data confirm that loop engineering is fully compatible with additional protein modification strategies and can be synergistically combined to optimize Acr inhibitory performance.

## Discussion

In this study, we systematically identified the loop region as a core and determinant functional module governing the inhibitory potency of the AcrIIC1 protein family. Compared with AcrIIC1_Nme1_, AcrIIC1_Boe_ exhibits substantially stronger Nme1Cas9 inhibitory activity in vitro. Consistent with previous structural reports, both AcrIIC1 homologs share a conserved binding mode toward the HNH domain and retain a canonical hydrogen-bonding network that shields the catalytic residues D587 and H588, which is essential for basal anti-CRISPR activity^21^. Nevertheless, mutational analysis of conserved catalytic-site residues failed to explain the superior inhibitory efficiency of AcrIIC1_Boe_, implying the existence of an additional, uncharacterized regulatory mechanism independent of HNH active-site shielding. This undiscovered mechanism is exactly the loop-mediated sgRNA loading interference firstly uncovered in our work, which well interprets the huge activity difference between the two homologous proteins. Further structural dynamics and functional assays uncovered a unique loop-mediated interference mechanism exclusive to AcrIIC1_Boe_. Molecular dynamics simulations revealed that the AcrIIC1_Boe_ loop creates prominent steric hindrance against the 5′ spacer region of sgRNA, increasing local structural flexibility of the Nme1Cas9-sgRNA complex^36^. This spatial obstruction is not a transient interaction but a persistent structural restriction, as validated by stopped-flow kinetic assays monitoring real-time R-loop formation dynamics^20^. Although loop-spacer interference does not drastically alter the global binding affinity between Nme1Cas9 and sgRNA, it effectively impedes stable R-loop assembly and disrupts the formation of functional Nme1Cas9-sgRNA-DNA ternary complexes. Consistently, limited proteolysis confirmed that the AcrIIC1_Boe_ loop increases the conformational flexibility of the Nme1Cas9-sgRNA complex^37^, directly linking loop-induced structural perturbation to impaired RNA binding, R-loop maturation, and target DNA recognition^38^. Site-directed mutagenesis of key loop residues further verified that the structural integrity of the loop is indispensable for this auxiliary regulatory function, establishing a solid structure–function relationship for this module.

Loop deletion mutagenesis across AcrIIC1 homologs further confirmed the essential and conserved role of the loop domain for both AcrIIC1_Nme1_ and AcrIIC1_Boe_. Loop removal completely abolished the inhibitory activity of both proteins. Distinct inhibitory modes exist between them: AcrIIC1_Nme1_ relies solely on HNH catalytic shielding for single-pathway inhibition, whereas AcrIIC1_Boe_ achieves potent suppression via two coupled and complementary inhibitory pathways, combining conserved HNH active-site occlusion and the newly identified sgRNA spacer interference. Even though the structure of AcrIIC1_Nme1_ has been resolved in prior high-profile studies, such divergent coupled regulatory modes have never been proposed^21–23^. Consistent with this mechanism, the naturally occurring inactive homolog AcrIIC1_Vei_, which possesses a conserved HNH-binding interface but lacks a stable functional loop interaction, exhibits no detectable inhibitory activity. Structural comparison revealed that the unstable loop-HNH interaction in AcrIIC1_Vei_ fails to support effective inhibition, further demonstrating that the loop domain serves as the fundamental functional core determining the antagonistic capacity of AcrIIC1 family proteins.

Loop-swapping experiments definitively validated the loop as a transferable functional module that dictates AcrIIC1 inhibitory potency, enabling artificial manipulation of anti-CRISPR activity. Grafting the incomplete loop of AcrIIC1_Nme1_ into AcrIIC1_Boe_ disrupted stable loop conformation, increased structural flexibility, and completely abolished inhibitory activity, despite retained catalytic-site hydrogen bond networks. Conversely, transplantation of the functional AcrIIC1_Boe_ loop into AcrIIC1_Nme1_ successfully recapitulated the spacer-interfering feature, significantly enhancing the inhibitory efficiency of AcrIIC1_Nme1_ to a level comparable to wild-type AcrIIC1_Boe_. Mammalian cell assays further confirmed that loop-mediated functional enhancement is robust in cellular environments^35^. Notably, loop engineering specifically potentiated inhibition toward Nme1Cas9 without broadening the inhibitory spectrum, indicating that loop-dependent regulation relies on the intrinsic interaction microenvironment of specific Cas9 orthologs. Furthermore, loop transplantation exhibited excellent compatibility with additional protein engineering strategies. The reported mCherry fusion acts as an independent modification module, while the loop region represents another novel functional unit^35^. The combined modification integrates three types of inhibitory effects within one protein, including inherent HNH suppression, mCherry-derived function and coupled dual-pathway inhibition from the transplanted loop. Such superimposed multiple mechanisms produce synergistic inhibitory effects, offering promising prospects to design anti-CRISPR variants with stronger inhibitory capacity and broader application scope in follow-up research.

In summary, this study reveals a coupled two-tiered inhibitory system of AcrIIC1 proteins controlled by a conserved and transferable loop module. While the loop maintains basal inhibitory function by stabilizing Acr–HNH interaction across homologs, the specialized loop conformation of AcrIIC1_Boe_ confers an additional sgRNA interference pathway that dramatically enhances anti-CRISPR potency through synergistic coupling with HNH shielding. Our findings uncover the structural basis of functional divergence within the AcrIIC1 family and provide new insights into the rational design and modular engineering of high-efficiency anti-CRISPR proteins for biotechnological applications.

## Materials and methods

### Protein expression and purification

Nme1Cas9 from *Neisseria meningitidis* was cloned and inserted into the expression vector pRSF-Duet-1 with a 6×His-Sumo tag at the N-terminus and a 6×His tag at the C-terminus. Nme1Cas9 (HNH domain) was subsequently cloned and inserted into the expression vector pGEX-6P-1. AcrIIC1 from *Brackiella oedipodis* was cloned and inserted into the expression vector pET-22b(+). The mutants were constructed using a site-directed mutagenesis kit. All proteins were overexpressed in *Escherichia coli* strain BL21 (DE3) (Genstar) cells and induced with 0.5 mM isopropyl-β-D-1-thiogalac-topyranoside (IPTG) at OD_600_=0.6 for 16 h at 16℃.

Cells that expressed Nme1Cas9 were lysed in a buffer containing 50 mM Tris-HCl (pH 7.5), 500 mM NaCl and 15% glycerol at 4℃. After centrifugation, the supernatant was incubated with Ni^2+^-Sepharose resin (GE Healthcare) and washed with a lysis buffer supplemented with 20 mM imidazole. The 6×His-Sumo tag was digested with His-tagged ubiquitin-like protein 1 (Ulp1) protease for 18 h at 4℃. The flow-through collection mixture was loaded onto an SP column (GE Healthcare) with reduced salt concentrations and eluted with a buffer containing 20 mM Tris-HCl (pH 7.5), 1 M NaCl, and 5 mM DTT. The resulting fractions that contained protein were purified using a Superdex 200 increase 10/300 gel filtration column and a buffer containing 20 mM Tris-HCl (pH 7.5), 150 mM NaCl and 5 mM DTT.

Cells expressing GST-HNH were lysed in 50 mM Tris-HCl (pH 7.5) and 500 mM NaCl buffer at 4℃. The HNH protein was further purified using glutathione Sepharose 4B affinity resin (Pharmacia) and washed with a lysis buffer. The GST tag was digested with commercial PreScission protease (Pharmacia) for 18 h at 4℃. Then, purification was performed using a procedure identical to that of Nme1Cas9.

Cells expressing 6×His-AcrIIC1 were lysed in 50 mM Tris-HCl (pH 7.5) and 500 mM NaCl buffer at ℃. The AcrIIC1 protein was further purified using Ni^2+^-Sepharose and Q HP columns (GE Healthcare) and eluted with a buffer containing 20 mM Tris-HCl (pH 7.5), 1 M NaCl and 5 mM DTT. Then, purification was performed using a procedure identical to that of Nme1Cas9.

The size exclusion binding assay was conducted by mixing the purified untagged Nme1Cas9 HNH domain with the purified untagged 1.2×AcrIIC1_Boe_ protein (or mutants) on ice and incubating for 1 h. Subsequently, the complexes were reprocessed on a Superdex 75 10/300 and analyzed on a 12% SDS-PAGE stained with Coomassie R250^39^.

### In vitro DNA cleavage assays

All reactions were performed in 1× reaction buffer (20 mM Tris-HCl, pH 7.5; 100 mM KCl; 5 mM MgCl_2_; 1 mM DTT; and 5% glycerol (v/v)). Nme1Cas9 complexes with sgRNA and four-fold Acr for 15 min at 0℃. The pUC19 target DNA plasmid (35- bp target DNA cloned into the pUC19 vector) was linearized by *Sca* I digestion before the cleavage reactions. Next, 300 ng of DNA substrate was added to 10 μL of reaction buffer, and the mixture was allowed to react for 15 min at 37℃. All reactions were stopped by adding 1 μL of 0.5 M EDTA (pH 8.0) and heated for 10 min at 70℃. The products were analyzed on 1% agarose, 1×TAE gels stained with ethidium bromide^39^.

### Pulldown assay

For the GST pull-down assays, Nme1Cas9 constructs (26 μg) were mixed with GST-tagged AcrIIC1_Boe_ (20 μg) in 600 μL of buffer containing 50 mM Tris-HCl (pH 7.5), 500 mM NaCl, and 15% glycerol and incubated with 40 μL GST Trap resin at 4 ℃ for 1 h. The resin was subsequently washed eight times with 800 μL of 50 mM Tris-HCl (pH 7.5), 500 mM NaCl, and 15% glycerol. Proteins were eluted by adding three resin volumes of 50 mM Tris-HCl pH 7.5, 500 mM NaCl, 15% glycerol, and 20 mM glutathione, resolved by sodium dodecyl sulfate polyacrylamide gel electrophoresis (SDS-PAGE) and quantified by Coomassie stain.

For the 6×His pull-down assays, AcrIIC1 mutants (26 ug) were mixed with 6 x His-tagged HNH_Nme1_ (20 μg) in 600 μL of buffer containing 50 mM Tris-HCl (pH 7.5), 500 mM NaCl, and 15% glycerol and incubated with 40 μL Ni Trap resin at 4 ℃ for 1 h. The resin was subsequently washed twelve times with 800 μL of 50 mM Tris-HCl (pH 7.5), 500 mM NaCl, 50 mM Imidazole and 15% glycerol. Proteins were eluted by adding three resin volumes of 50 mM Tris-HCl pH 7.5, 500 mM NaCl, 15% glycerol, and 300 mM Imidazole, resolved by sodium dodecyl sulfate polyacrylamide gel electrophoresis (SDS-PAGE) and quantified by Coomassie stain.

### Limited proteolysis of Nme1Cas9-AcrIIC1 complex

Limited α-chymotrypsin proteolysis assays were performed at 25 ℃ for different times (0, 10, 30, 60 min) using proteolysis buffer 20 mM Tris pH 7.5, 300 mM NaCl. The same amount (80 μg) of purified Nme1Cas9 was used to construct complex Nme1Cas9-sgRNA (1:1.1), Nme1Cas9-AcrIIC1 (1:4), Nme1Cas9-AcrIIC1-sgRNA (1:4:1.1). The reactions were stopped by adding 5×SDS loading buffer and quenched for 10 min at 100 °C. Samples were analyzed on a 12% SDS polyacrylamide gel with Tris-Gly running buffer and stained with Coomassie Brilliant Blue R-250^40^.

### Crystallization, data collection and structure determination

All crystals were grown using the microbatch-under-oil method unless otherwise specified. AcrIIC1_Boe_ was crystallized at 16℃ by mixing 1 µL of protein (5 mg mL^-1^) with 1 µL of crystallization buffer, which contained 800 mM sodium phosphate monobasic, 1200 mM potassium phosphate dibasic and 100 mM sodium acetate/acetic acid (pH 4.5). The crystals were cryoprotected by Parabar 10312 (previously known as paratone oil; Hampton Research, USA). X-ray diffraction data were collected on beamline BL18U1 at the Shanghai Synchrotron Radiation Facility at 100 K and a wavelength of 0.97776 Å. Data integration and scaling were performed using HKL3000^41^. The structure was determined by molecular replacement, where the structure of AcrIIC1_Nme_ (PDB ID: 5VGB) was used as the search model.

The AcrIIC1_Boe_-HNH domain of Nme1Cas9 was crystallized at 16°C by mixing 1 µL of protein (5 mg/mL) with 1 µL of crystallization buffer containing 200 nM ammonium sulfate, 100 mM sodium acetate (pH 4.5), and 22% (w/v) PEG4000. X-ray diffraction data were collected on beamline BL18U1 at the Shanghai Synchrotron Radiation Facility. The structure was determined by molecular replacement, where the structure of AcrIIC1nme (PDB ID: 5VGB) was used as the search model.

The AcrIIC1_Vei_-HNH domain of Nme1Cas9 was crystallized at 16°C by mixing 1 µL of protein (5 mg mL^-1^) with 1 µL of crystallization buffer containing 30% (v/v) 2-Propanol, 100 mM TRIS /Hydrochloric Acid (pH 8.5), and 30% (w/v) PEG 3350. X-ray diffraction data were collected on beamline BL02U1 at the Shanghai Synchrotron Radiation Facility. The structure was determined by molecular replacement, where the structure of AcrIIC1nme (PDB ID: 5VGB) was used as the search model.

The AcrIIC1_Nme1ST_-HNH domain of Nme1Cas9 was crystallized at 16°C by mixing 1 µL of protein (5 mg mL^-1^) with 1 µL of crystallization buffer containing 0.1 M sodium citrate tribasic dihydrate (pH 5.0) and 10% (w/v) PEG 6,000. X-ray diffraction data were collected on beamline BL02U1 at the Shanghai Synchrotron Radiation Facility. The structure was determined by molecular replacement, where the structure of AcrIIC1nme (PDB ID: 5VGB) was used as the search model.

### Gel shift assays

Binding reactions were conducted in Binding Buffer (20 mM Tris-HCl, pH 7.5, 150 mM KCl, 5 mM EDTA, 5 mM MgCl_2_, 1 mM DTT, 5% (v/v) glycerol). Nme1Cas9 and sgRNA were incubated first for 5 min at 37℃ to allow for guide binding. Next the Nme1Cas9-sgRNA complex was diluted to the indicated concentration and a constant amount of 20 mM Acr was added to each sample and allowed to incubate for 10 min at room temperature. 100 ng Fam-labeled DNA target was then added. The binding reaction was incubated at 37℃ for 30 min at room temperature. Samples were analyzed by 6% polyacrylamide/0.5×TBE gel electrophoresis. The results were characterized by a gel imaging system. Assays were conducted in triplicate with representative gels shown^42^.

### Bioinformatics analysis

The BLASTp program was used to compare the percent query cover and identity among type-II Cas9 orthologs. The protein sequences of representative type-II-C Cas9 orthologs were aligned via Jalview (https://www.jalview.org/). The structure of the HNH domains from Cas9 orthologs (available) was determined using PyMol (https://pymol.org/2/).

### Molecular Dynamics Simulations

The 100-ns all-atom simulations were performed on Gromacs 2020.6 with three independent trajectories^43,44^. First, the X-ray structural model of Nme1Cas9 (PDB ID: 6JDQ) was relaxed and equilibrated without sgRNA. Then, flexible docking was performed to obtain complexes of Nme1Cas9 and full-length sgRNA^45^, followed by 100-ns simulations using the CHARMM36 force field^46^. The system was solvated with TIP3P water and ions (0.10 M NaCl and 0.05 M MgCl_2_) in a 4458.23-nm^3^ box (451983 atoms in total). The Nose-Hoover thermostat (303.15 K) and Parrinello-Rahman isotropic NPT ensemble were used with h-bond LINCS constraints. After the system had reached equilibrium, snapshots of each trajectory were clustered and analyzed for binding conformations.

### Protein structure prediction using AlphaFold 3

Structure predictions for seven representative AcrIIC1 family members were performed using the AlphaFold 3 Server (https://alphafoldserver.com)^47^. The full-length amino acid sequences were submitted with default parameters. For each predicted structure, the highest-ranking model (ranked_0) based on the predicted template modeling (pTM) score was selected for structural analysis and figure preparation. Structural figures were generated using PyMOL.

### Binding affinity

The binding affinity assays were detected using BLI (Octet® R8, Sartorius). The complexed Nme1Cas9 and Acr proteins were incubated with a buffer containing 20 mM Tris-HCl (pH 7.5), 150 mM NaCl, and 5 mM DTT for 30 min at 0℃. The complexed proteins were immobilized on NTA biosensors (Sartorius) and incubated with sgRNA. The binding affinities were calculated and analyzed using Octet® BLI and Origin 2023b^20^.

### TIDE sequencing

Genomic target sites relevant for TIDE experiments are listed in **Table S2 and S3**. For transfection-based experiments, CHO cells were seeded in 6-well plates at a density of 500,000 cells per well. Transfections were conducted with Lipofectamine 3000 using 7.5 μL of Lipofectamine reagent, 5.0 μL of P3000 and 2.4 μg of total DNA per well. Cells were cotransfected with 1.6 μg of Acr vector, 0.4 μg of Nme1Cas9 vector and 0.4 μg of sgRNA vector^35,48^.

3 days after transfection, genomic DNA was extracted from the cells using a genomic DNA extraction kit (TianGen). PCR amplification of the CRISPR-Cas9 targeted genomic locus was then performed using a PCR enzyme (Genestar) and appropriate primers. The indel frequency was evaluated by TIDE sequencing (https://tide.deskgen.com/).

### Stopped-flow method

Perform the stopped flow measurement by rapidly mixing 500 nM dNme1Cas9-sgRNA complex or dNme1Cas9-AcrIIC1-sgRNA complex (Syringe 1) with 500 nM labeled DNA duplex (Syringe 2) in the reaction buffer.

Since the emitted light of Cy-3 can serve as the excitation light for Cy-5, as the DNA double helix unfolds and the distance increases, the emission intensity of Cy-5 will weaken. Therefore, we can measure the rate of R-loop formation by using the flow spectrometer (Bio-Logic) to record the changes in fluorescence intensity. Excite the fluorophore at 550 nm and monitor the fluorescence change at 670 nm^28^.

## Reporting summary

Further information on research design is available in the Nature Portfolio Reporting Summary linked to this article.

## Data availability

The PDB accession codes for the coordinates of AcrIIC1_Boe_, HNH_Nme1_-AcrIIC1_Boe_, HNH_Nme1_-AcrIIC1_Vei_, HNH_Nme1_-AcrIIC1_Nme1ST_, and HNH_Nme1_-AcrIIC1_BoeST_ are 9JB6, 9J1Y, 9J2G, 9J1Z, and 9WN6 respectively.

## Author contributions

Z.W., Y.X. and H.Y. conceived the project; Z.W., Y.X. and S.W. designed the experiments; S.W., Z.Z., W.L., Q.W., X.L., Chang Liu. and Chenyu Luan. cloned, expressed, purified and crystallized protein; S.W., Y.X. and Q.W. collected the diffraction data; Y.X. and S.W. solved the crystal structure; S.W. and Z.Z. analyzed and discussed the data; C.Z. carried out molecular dynamic simulation experiments and data analysis; Z.W., S.W., Y.X., H.Y. and C.Z. wrote the manuscript.

## Declaration of competing interest

The authors declare no competing interests.

## Acknowledgments

We thank the staff from beamline BL10U2, BL19U1, BL18U1 and BL02U1 at Shanghai Synchrotron Radiation Facility (SSRF) for the strong support of our data collection and processing. This work was supported by grants from the Tianjin Science and Technology Project (No. 25ZXDFQY00120) to Z.W., the National Natural Science Foundation of China (No. 31970048) to Z.W., the National Natural Science Foundation of China (NO. 82202518) to Y.X.

## Notes

### Competing Interest Statement

The authors have declared no competing interest.

