## Supplemental Information for "Structural basis for loop-dependent inhibition by AcrIIC1 anti-CRISPR proteins"

**This PDF file includes:**

Figures S1 to S20

Tables S1 to S3

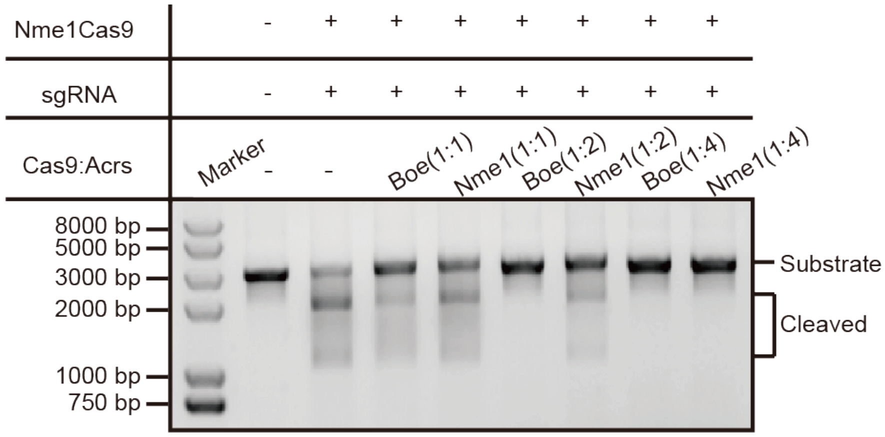

**Figure S1. DNA cleavage assays performed by Nme1Cas9 in the presence of AcrIIC1$_{Boe}$ and AcrIIC1$_{Nme1}$ at various ratios**

In order to compare the antagonistic efficiency of AcrIIC1$_{Boe}$ and AcrIIC1$_{Nme1}$, AcrIIC1$_{Boe}$ and AcrIIC1$_{Nme1}$ with various molar ratios were used to inhibit the cleavage of target DNA by Nme1Cas9 in vitro.

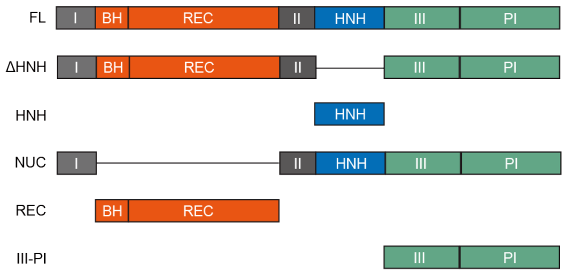

**Figure S2. Schematic diagram of the full-length and truncated domains of Nme1Cas9.**

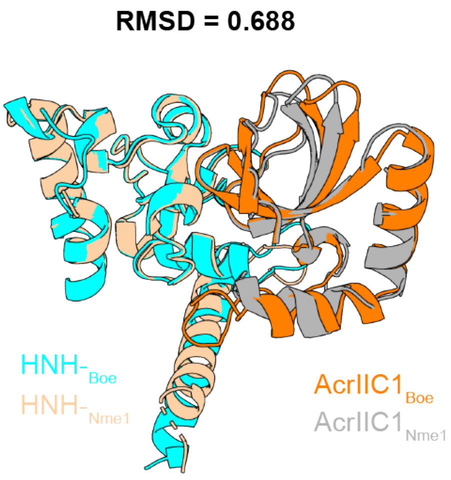

**Figure S3. Structural alignment of the AcrIIC1$_{Boe}$-HNH and AcrIIC1$_{Nme1}$-HNH complexes.** Structural superposition of the AcrIIC1$_{Boe}$-HNH complex and the AcrIIC1$_{Nme1}$-HNH complex (PDB: 5VGB) shown in cartoon representation. AcrIIC1$_{Boe}$ is colored orange, and the HNH domain from the AcrIIC1$_{Boe}$ complex is colored cyan. AcrIIC1$_{Nme1}$ is colored gray, and the HNH domain from the AcrIIC1$_{Nme1}$ complex is colored wheat. The alignment yields an RMSD of 0.688 Å over the core fold.

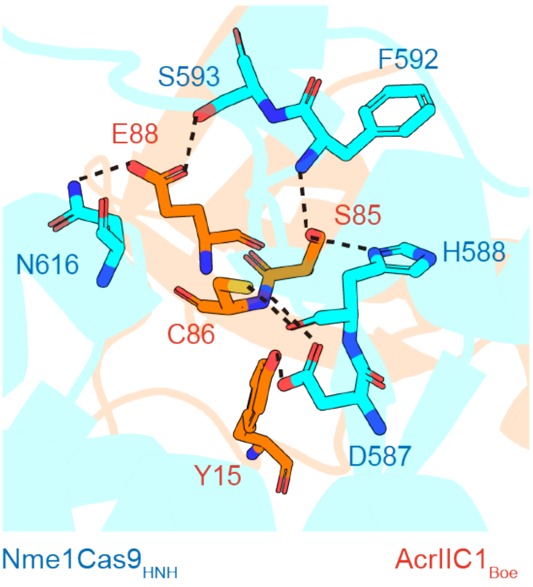

**Figure S4 Active site residues of the HNH domain, which are shielded by AcrIIC1$_{Boe}$ via hydrogen bonding.**

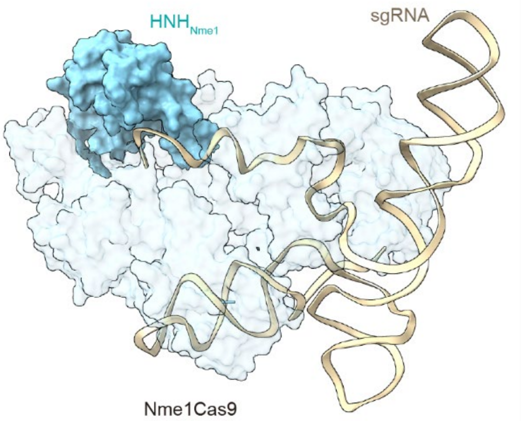

**Figure S5. Molecular dynamics simulation of the Nme1Cas9-sgRNA complex**
The HNH domain is shown in cyan, and sgRNA in wheat.

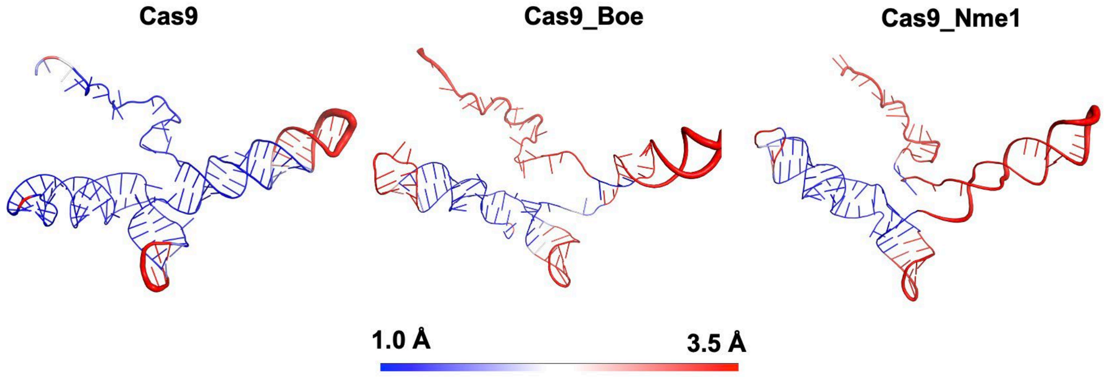

**Figure S6. The conformational fluctuations of sgRNA in different bound state.**
Red means more flexible; Blue means less flexible.

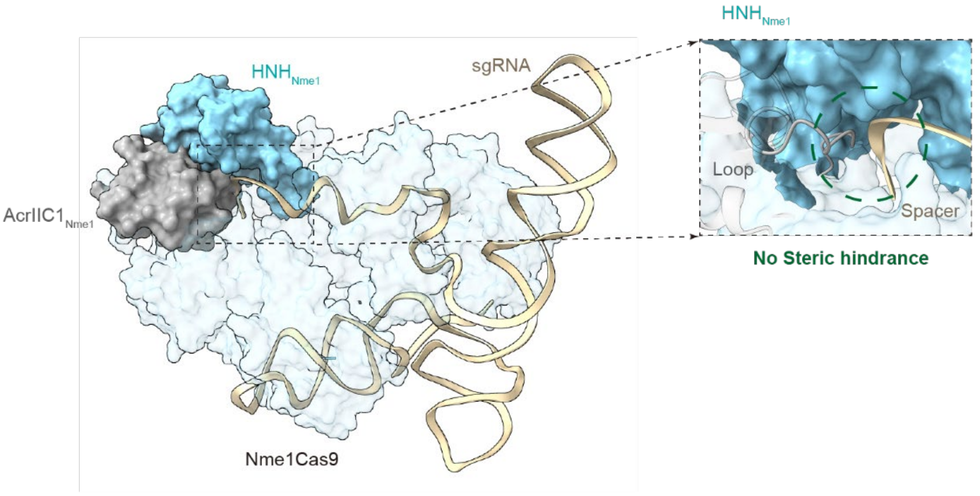

**Figure S7. Molecular dynamics simulation of the Nme1Cas9-sgRNA complex in the presence of AcrIIC1$_{Nme1}$.**
AcrIIC1$_{Nme1}$ is shown in gray, the HNH domain in cyan, and sgRNA in wheat.

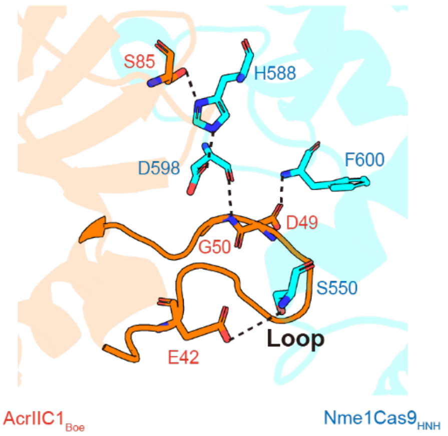

**Figure S8. Hydrogen-bond interactions between the AcrIIC1$_{Boe}$ loop and the HNH domain observed in the AcrIIC1$_{Boe}$-HNH crystal structure.**

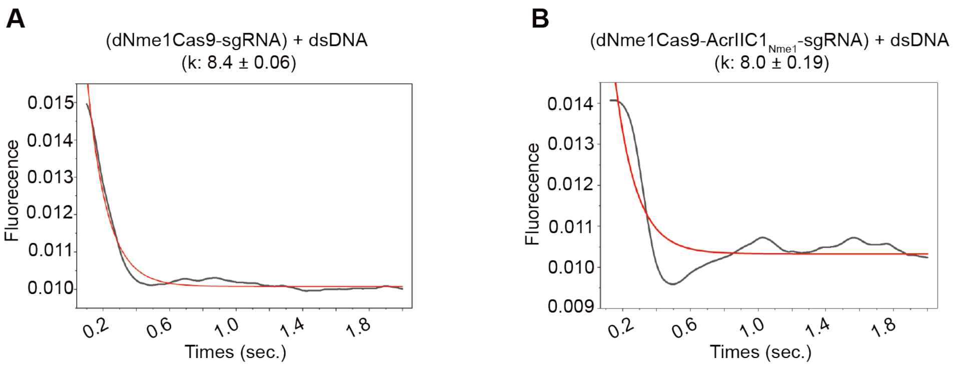

**Figure S9 Analyzing and fitting the data of R-loop formation**

**A,** Analysis of the rate of R-loop formation in the absence of AcrIIC1 protein **B,** In the presence of AcrIIC1$_{Nme1}$, the analysis of the rate of R-loop formation.

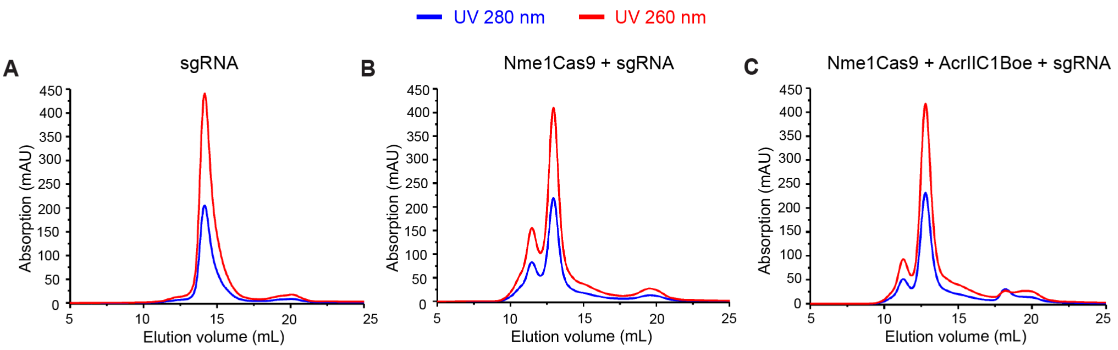

**Figure S10. Size exclusion chromatography of sgRNA, Nme1Cas9-sgRNA and Nme1Cas9-AcrIIC1$_{Boe}$-sgRNA**

**A,** Size exclusion chromatography of sgRNA**. B,** Size exclusion chromatography of Nme1Cas9-sgRNA complex. **C,** Size exclusion chromatography of Nme1Cas9-AcrIIC1$_{Boe}$-sgRNA complex.

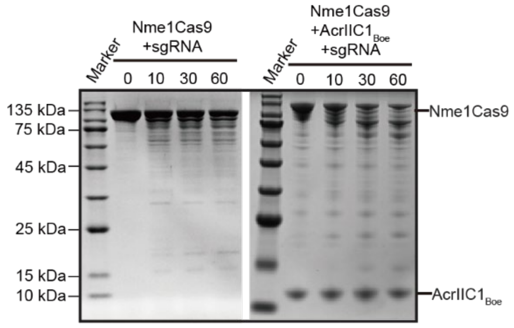

**Figure S11. Restriction enzyme digestion results of the Nme1Cas9-sgRNA and Nme1Cas9-AcrIIC1$_{Boe}$-sgRNA complexes.**

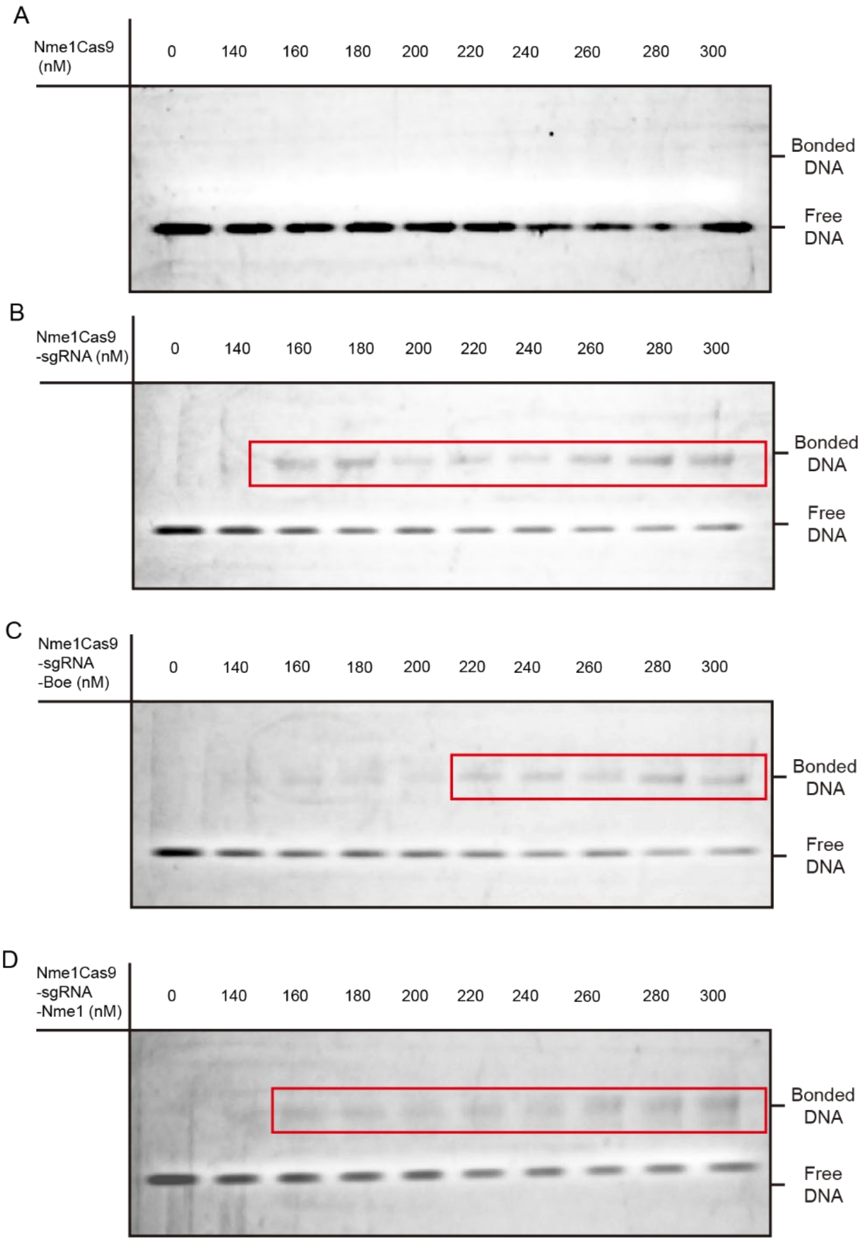

**Figure S12. Analysis of the binding of Nme1Cas9 to the labeled DNA in the presence or absence of AcrIIC1 on a non-denaturing gel.**

**A,** The binding of Nme1Cas9 to the labeled DNA. **B,** The binding of Nme1Cas9-sgRNA to the labeled DNA. **C,** The binding of Nme1Cas9-sgRNA-AcrIIC1$_{Boe}$ to the labeled DNA. **D,** The binding of Nme1Cas9-sgRNA-AcrIIC1$_{Nme1}$ to the labeled DNA.

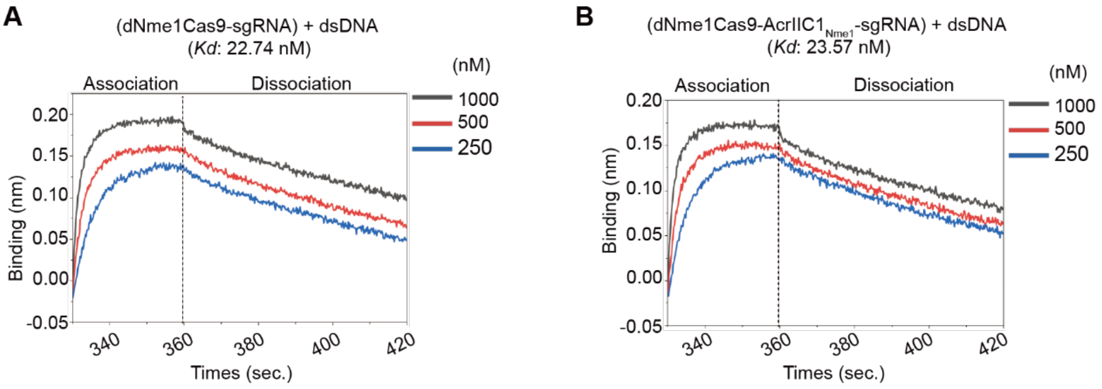

**Figure S13 Binding affinity measurements between dNme1Cas9-sgRNA and dsDNA.**
**A,** Binding affinity measurements between dNme1Cas9-sgRNA and dsDNA. **B,** Binding affinity measurements between dNme1Cas9-sgRNA and dsDNA in the presence of AcrIIC1$_{Nme1}$

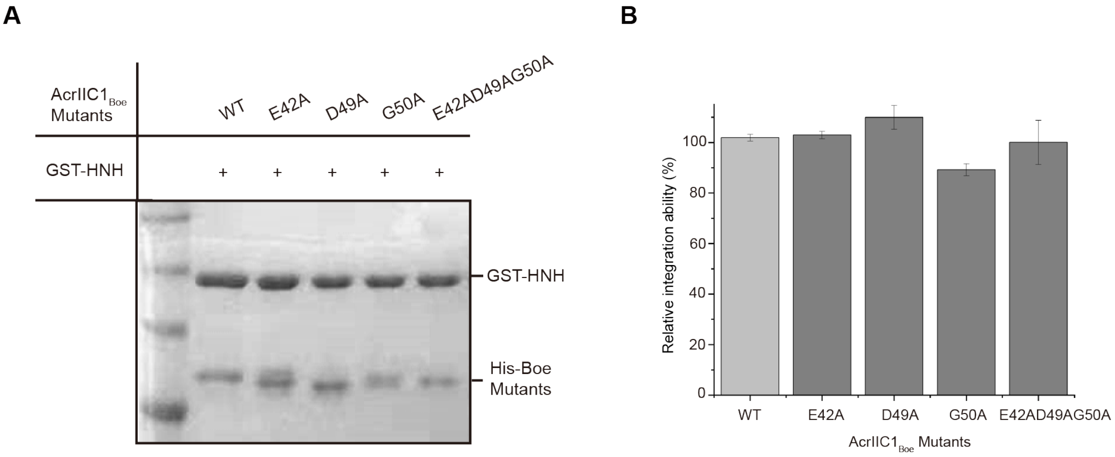

**Figure S14. Pull-down results of WT AcrIIC1Boe and the indicated mutants in the loop region with the HNH domain.**

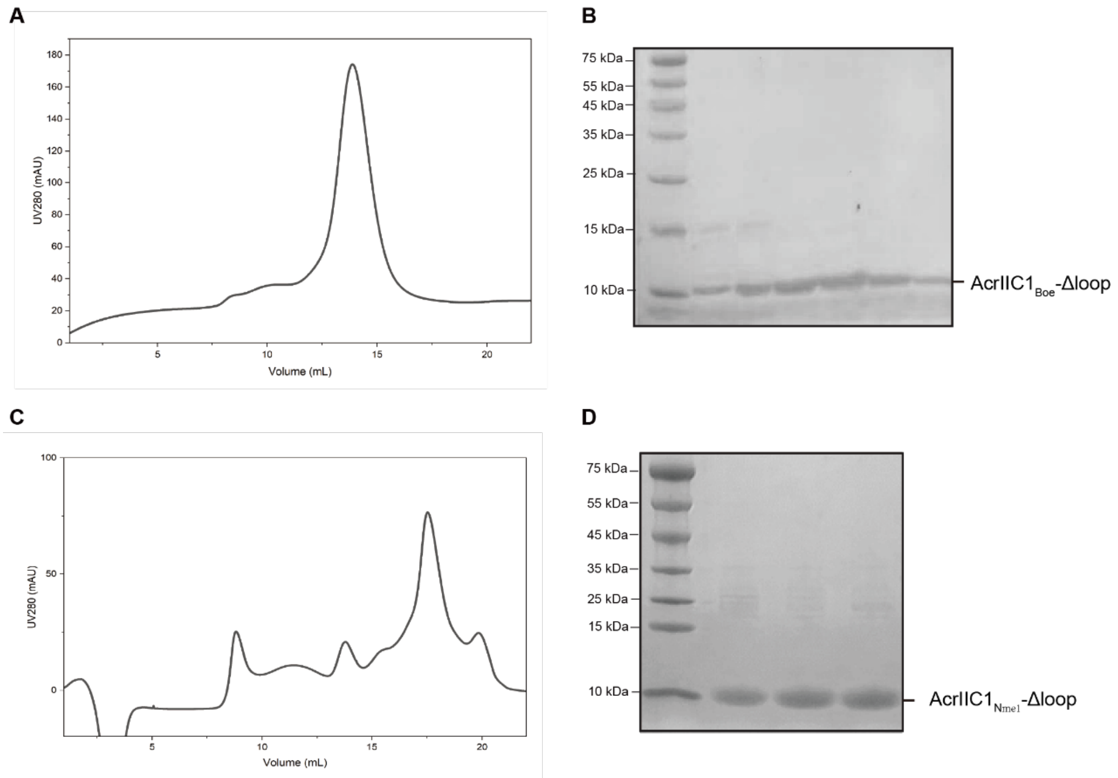

**Figure S15. Size exclusion chromatography and SDS-PAGE of AcrIIC1$_{Boe}$-Δloop and AcrIIC1$_{Nme1}$-Δloop**

**A,** The gel filtration chromatography peak map of AcrIIC1$_{Boe}$-Δloop; **B,** The SDS-PAGE corresponding to the gel filtration chromatography of AcrIIC1$_{Boe}$-Δloop; **C,** The gel filtration chromatography peak map of AcrIIC1$_{Nme1}$-Δloop; **D,** The SDS-PAGE corresponding to the gel filtration chromatography of AcrIIC1$_{Nme1}$-Δloop.

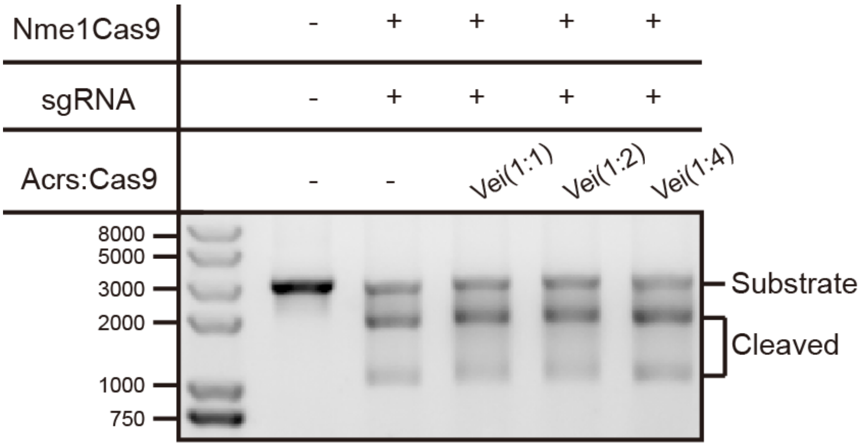

**Figure S16. DNA cleavage assays performed by Nme1Cas9 in the presence of AcrIIC1$_{Vei}$ at various ratios**

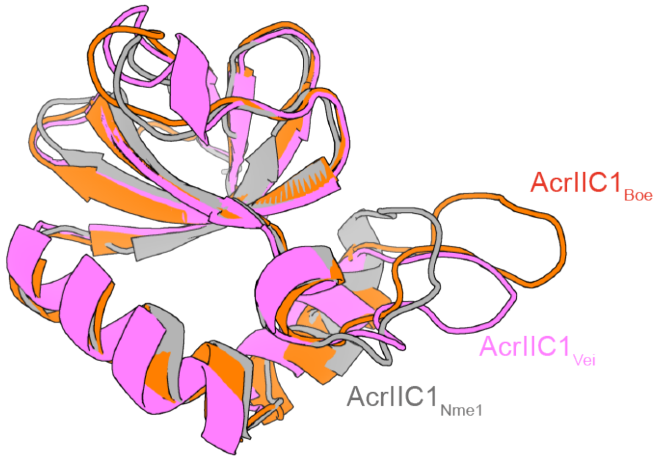

**Figure S17. Structural alignment of the AcrIIC1$_{Boe}$, AcrIIC1$_{Nme1}$ and AcrIIC1$_{Vei}$.**

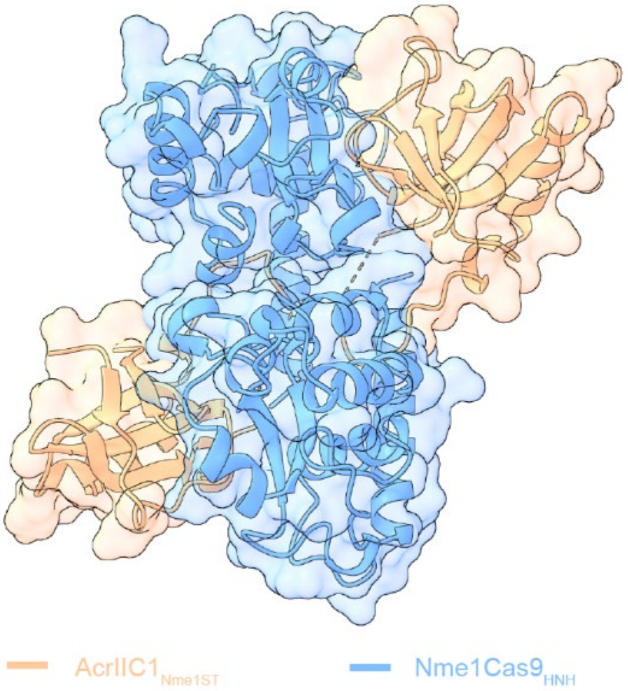

**Figure S18. The overall crystal structure of Nme1Cas9 HNH domain and AcrIIC1$_{Nme1ST}$ complex**

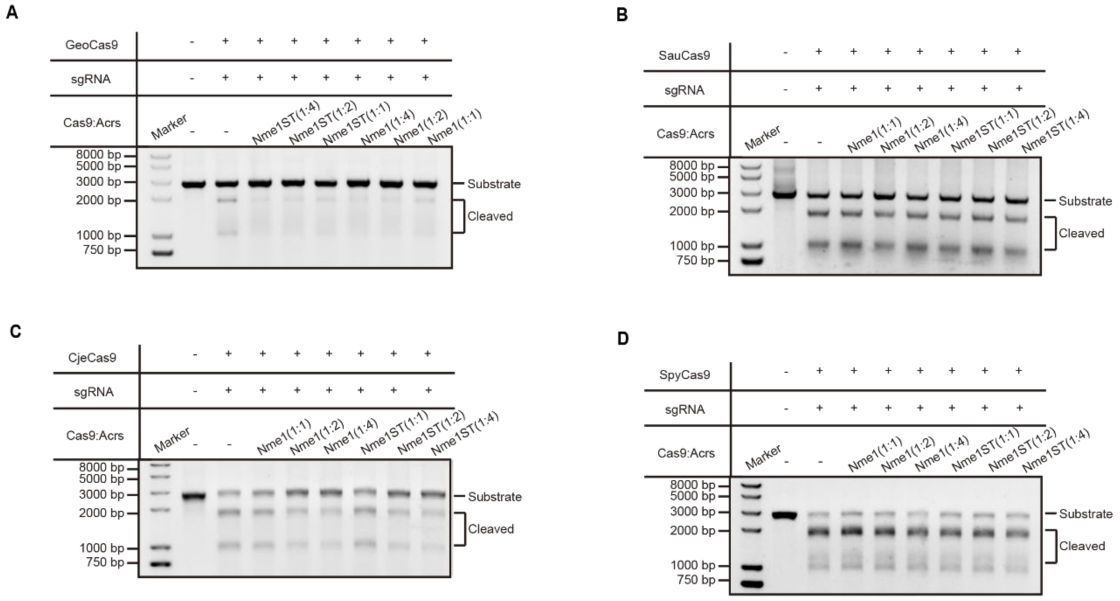

**Figure S19. In vitro cleavage inhibition experiments of AcrIIC1$_{Nme1}$ and AcrIIC1$_{Nme1ST}$ on II-A or II-C type Cas9.**

**A,** In vitro cleavage inhibition experiment of GeoCas9 by AcrIIC1$_{Nme1}$ and AcrIIC1$_{Nme1ST}$. **B,** In vitro cleavage inhibition experiment of SauCas9 by AcrIIC1$_{Nme1}$ and AcrIIC1$_{Nme1ST}$. **C,** In vitro cleavage inhibition experiment of CjeCas9 by AcrIIC1$_{Nme1}$ and AcrIIC1$_{Nme1ST}$. **D,** In vitro cleavage inhibition experiment of SpyCas9 by AcrIIC1$_{Nme1}$ and AcrIIC1$_{Nme1ST}$.

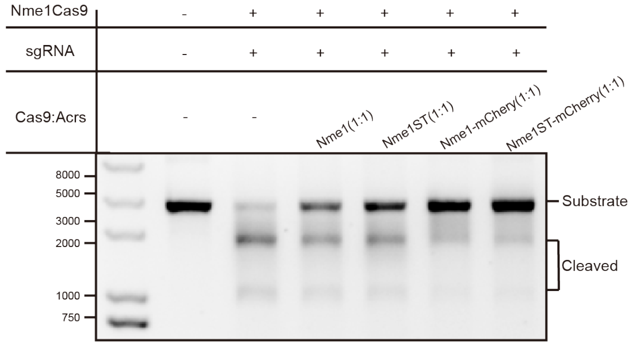

**Figure S20. DNA cleavage assays conducted by Nme1Cas9 in the presence of AcrIIC1$_{Nme1}$, AcrIIC1$_{Nme1ST}$, AcrIIC1$_{Nme1}$-mCherry and AcrIIC1$_{Nme1ST}$-mCherry.**

**Table S1. X-ray data collection and refinement statistics.**

| | AcrIIC1$_{Boe}$ | HNH$_{Nme1}$-AcrIIC1$_{Boe}$ | HNH$_{Nme1}$-AcrIIC1$_{Vei}$ | HNH$_{Nme1}$-AcrIIC1$_{Nme1ST}$ | HNH$_{Nme1}$-AcrIIC1$_{BoeST}$ |
|---|---|---|---|---|---|
| Space group | $I2_13$ | $P3_1$ | $P65$ | $P1$ | $P12_11$ |
| Cell dimensions | | | | | |

| | | | | | |
|---|---|---|---|---|---|
| *a, b, c* (Å) | 92.88, 92.88, 92.88 | 61.75, 61.75, 50.03 | 76.24, 76.24, 89.42 | 45.96, 47.63, 61.72 | 45.47, 110.21, 46.99 |
| α, β, γ (°) | 90.0, 90.0, 90.0 | 90.0, 90.0, 120.0 | 90.00, 90.00, 120.00 | 107.44, 97.49, 115.25 | 90.00, 116.16, 90.00 |
| Wavelength (Å) | 0.9778 | 0.9785 | 0.9792 | 0.9792 | 0.9792 |
| Resolution limit (Å) | 50.00-2.40 (2.49-2.40)* | 50.00-1.80 (1.85-1.80)* | 50.00-2.10 (2.15-2.10)* | 50.00- 1.91 (1.98-1.91)* | 50-1.70 (1.76-1.70)* |
| $R_{merge}$ (%) | 15.50 (158.00) | 14.60 (158.00) | 21.70 (134.20) | 7.90 (27.7) | 12.00 (111.10) |
| $I / \sigma I$ | 30.53 (4.79) | 17.66 (2.94) | 15.30 (5.10) | 15.62 (5.88) | |
| Completeness (%) | 99.57 (99.43) | 99.05 (92.12) | 99.95 (100.00) | 96.37 (82.91) | 91.48 (81.94) |
| Redundancy | 38.70 (35.90) | 10.10 (8.50) | 20.70 (19.30) | 3.40 (2.70) | 2.90 (2.60) |
| **Refinement** | | | | | |
| Resolution range (Å) | 50.00-2.40 | 50.00-1.80 | 50.00-2.10 | 50.00- 1.90 | 50.00-1.70 |
| No. reflections | 5333 (522) | 19911 (1860) | 17757 (1760) | 32129 (2750) | 41742 (3475) |
| $R_{work}$ / $R_{free}$ | 0.203/0.252 | 0.160/0.188 | 0.174/0.192 | 0.183/0.215 | 0.166/0.215 |
| NO. atoms | | | | | |
| Protein | 725 | 2131 | 1933 | 3729 | 3673 |
| Ligand/ion | 0 | 0 | 0 | 0 | 0 |
| Water | 27 | 233 | 131 | 279 | 540 |
| *B*-factors | | | | | |
| Protein | 38.60 | 23.30 | 37.84 | 25.40 | 21.54 |
| Ligand/ion | / | / | / | / | / |
| Water | 36.18 | 31.34 | 45.45 | 33.20 | 32.58 |
| Bond lengths (Å) | 0.009 | 0.009 | 0.004 | 0.008 | 0.008 |
| Bond Angles (°) | 0.92 | 0.94 | 0.60 | 0.90 | 0.98 |
| Ramachandran plot | | | | | |
| Favored（%） | 96.63 | 98.69 | 97.32 | 97.45 | 98.63 |
| Allowed（%） | 3.37 | 1.31 | 2.68 | 2.55 | 1.37 |

| | | | | | |
|---|---|---|---|---|---|
| Outliers（%） | 0.00 | 0.00 | 0.00 | 0.00 | 0.00 |

*Values in parentheses are for highest-resolution shell.

**Table S2. Genomic target sites.** The sgRNA-complementary part is underlined. The PAM is shown in bold.

| Locus | Target sequence (5' to 3') |
|---|---|
| AAVS1 | ACCCCACAGTGGGGCCACTAGGGACAG**GATT** |

**Table S3. Primers for genomic PCRs.** Fw, forward primer; rv, reverse primer.

| Locus | Direction | Sequence (5' to 3') |
|---|---|---|
| AAVS1 | fw | TGCTTTCTTTGCCTGGACAC |
| | rv | CCTCTCTGGCTCCATCGTAA |
